# Functional dissection of basal ganglia inhibitory input onto SNc dopaminergic neurons

**DOI:** 10.1101/856617

**Authors:** RC Evans, EL Twedell, M Zhu, J Ascencio, R Zhang, ZM Khaliq

## Abstract

Substania nigra (SNc) dopaminergic neurons show a pause-rebound firing pattern in response to aversive events. Because these neurons integrate information from predominately inhibitory brain areas, it is important to determine which inputs functionally inhibit the dopamine neurons and whether this pause-rebound firing pattern can be produced by a solely inhibitory input. Here, we functionally map genetically-defined inhibitory projections from the dorsal striatum (striosome and matrix) and globus pallidus (GPe; parvalbumin and Lhx6) onto SNc neurons. We find that GPe and striosomal inputs both pause firing in SNc neurons, but rebound firing only occurs after inhibition from striosomes. Indeed, we find that striosomes are synaptically optimized to produce rebound and preferentially inhibit a subpopulation of ventral, intrinsically rebound-ready SNc dopaminergic neurons on their reticulata dendrites. Therefore, we describe a self-contained dendrite-specific striatonigral circuit that can produce pause-rebound firing in the absence of excitatory input.

## Introduction

Midbrain dopaminergic (SNc) neurons are activated by rewarding events (Schultz et al., 1997) and inhibited by aversive events (Matsumoto and Hikosaka, 2009). A subset of dopaminergic neurons exhibit rebound activity at the termination of an aversive stimulus (Brischoux et al., 2009; Fiorillo et al., 2013a; Wang and Tsien, 2011), which may serve as a safety or learning signal (Oleson et al., 2012; Lee et al., 2016; Schultz, 2019). Pause-rebound firing in SNc neurons likely involves inhibitory input which arrives from a variety of brain nuclei (Lerner et al., 2015; Menegas et al., 2015; Watabe-Uchida et al., 2012) and comprises up to 70% of synapses formed onto dopaminergic neurons (Henny et al., 2012). Therefore, to understand the pause-rebound firing pattern, it is critical to determine the functional impact of inhibitory inputs on excitability of SNc dopaminergic neurons.

Within the basal ganglia, the dorsal striatum and the external globus pallidus (GPe) are prominent sources of inhibition onto SNc neurons. However, a functional understanding of their direct connections to the dopamine neurons is complicated by the multiple cell types located within GPe and the striatum. For example, the GPe contains distinct subpopulations (Mallet et al., 2012; Mastro et al., 2014; Hernández et al., 2015), including parvalbumin- and Lhx6-positive neurons that project to substantia nigra and selective stimulation of these populations can differentially rescue Parkinsonian motor deficits (Mastro et al., 2017).

The dorsal striatum can be divided into neurochemically distinct subcompartments called striosomes (patches) and matrix (Graybiel et al., 1981; Gerfen et al., 1987). Axons from striosomes form dense bundles around the dendrites of SNc dopamine neurons termed ‘striosome-dendron bouquets’ (Crittenden et al., 2016), suggesting that the striosomes are a strong source of striatal inhibition onto the dopamine neurons. The first monosynaptic tracing study examining inputs onto SNc neurons showed a predominate input from striosomes (Watabe-Uchida et al., 2012). However, a more recent tracing study showed a higher number of labeled matrix neurons than striosome neurons (Smith et al., 2016), suggesting that the matrix is the stronger source of striatal inhibition onto SNc dopaminergic neurons. Importantly, these viral tracing methods provide an estimate of the number of pre-synaptically connected cells, but it is unclear how cell number relates to connection strength. These conflicting anatomical studies make it clear that understanding the source of inhibition onto the SNc dopamine neurons will require the functional testing of striosome and matrix inputs.

Functionally testing the strength and characteristics of inhibitory inputs onto SNc neurons will allow us to determine which specific inhibitory sources can induce rebound activity. Interestingly, we have previously shown that only certain subpopulations of dopamine neurons contain the intrinsic mechanisms necessary to rebound from inhibition, while other subpopulations lack this feature (Evans et al., 2017; Tarfa et al., 2017). This suggests that the pause-rebound firing pattern would be limited to a distinct population of dopamine neurons. However, it is unclear whether these SNc subpopulations are embedded in the same inhibitory circuit, or whether the inhibitory inputs to SNc neurons are subpopulation specific. Furthermore, the intrinsic rebound mechanisms of SNc neurons are located on dendrites and require hyperpolarization to be recruited. Therefore, the synaptic characteristics and subcellular location of each inhibitory input will play a critical role in their ability to generate dopamine rebound.

Here, we perform a functional dissection and dendritic mapping of genetically-defined sources of inhibition onto SNc neurons. We find that the dorsal striatum innervates the SNc primarily through striosomes and preferentially inhibits a subpopulation of rebound-ready SNc dopamine neurons. Therefore, we reveal an inhibitory striatonigral circuit which is both synaptically and intrinsically optimized to induce dopamine rebound.

## Results

### Functional test of genetically-defined inhibitory inputs to SNc dopamine neurons

To test the functional strength of striosome and matrix inputs to the SNc dopamine neurons, we injected AAV1-FLEX-hSyn-CoChR-GFP into the dorsal striatum of Pdyn-IRES-Cre mice to infect striosome projections and calbindin-IRES-Cre mice to infect matrix projections (Figure 1A). Imaging these axons in cleared brain slices stained for tyrosine hydroxylase (TH), we saw that the striosomal axons form clear axon bundles around the ventrally-projecting dopamine neuron dendrites, while axons from the striatal matrix fill in the SNr relatively evenly (Figure 1B), in agreement with previous work (Crittenden et al., 2016).

**Figure 1.**
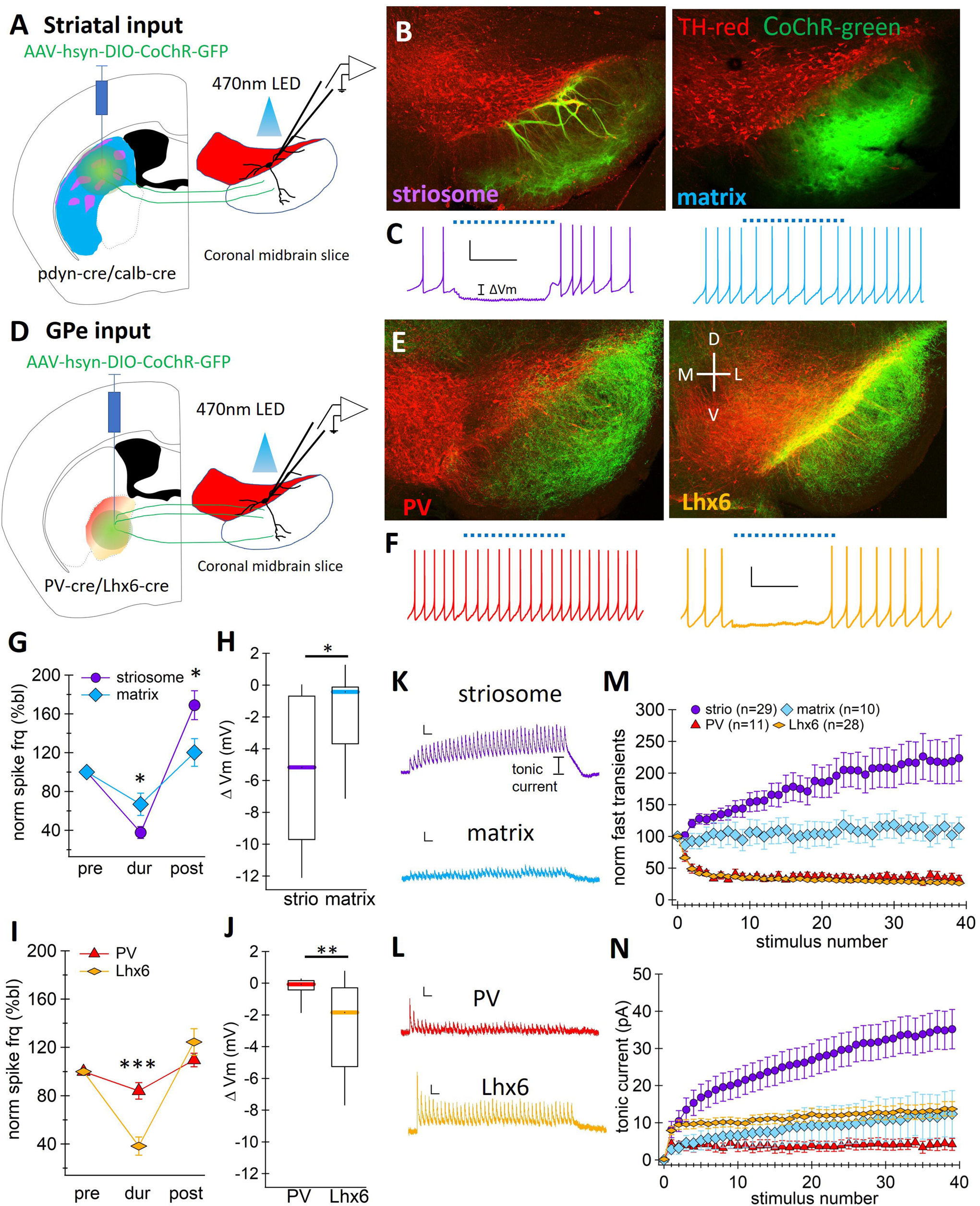
Genetically-defined basal ganglia inputs from dorsal striatum and globus pallidus differentially inhibit SNc dopamine neurons. **A.** Schematic of striatal injection site. **B.** Image of coronal brain slices stained for tyrosine hydroxylase (red) with striatal axons (green) from the striosomes (left) and matrix (right). **C.** Example traces from SNc dopamine neurons in response to optogenetic activation of striosomal (left) and matrix (right) axons. Scale bars: 20 mV, 1 second. **D-F.** Same as A-C, but for parvalbumin (PV, left) and Lhx6 (right) globus pallidus (GPe) inputs to SNc dopamine neurons. **G.** Summary of normalized action potential firing frequency before (pre), during (dur), and after (post) optogenetic activation of striatal fibers. **H.** Box plot of average membrane hyperpolarization in response to optogenetic activation of striosomal (strio) and matrix inputs **I-J.** Same as G-H, but for PV and Lhx6 GPe input. **K.** Voltage-clamp traces of inhibitory synaptic currents in SNc dopamine neurons in response to optogenetic activation of inputs from striosomes (top) and matrix (bottom). Scale bars: 20 pA, 100 ms. **L.** same as K, but for GPe subpopulations. **M.** Analysis of short-term plasticity (normalized transient current amplitude) during stimulus train **N.** Average amplitude of tonic current component over the course of the 20 Hz, 2-second optogenetic stimulation train. *p<0.05; **p<0.01, ***p<0.001

We compared the effect of inhibition from either striosomal or matrix inputs on SNc neuron firing by applying 20 Hz light stimulation (470 nm LED) for 2 seconds. We found that activation of striosomal inputs resulted in stronger inhibition of tonic firing and more effective hyperpolarization of SNc dopamine neurons than activation of matrix inputs (normalized spike frequency during inhibition as percent baseline; striosomes 37.6 ± 6.27% n=56, matrix 66.9 ± 11.4% n=15, p=0.035, Wilcoxon Rank Test; avg Vm during inhibition relative to baseline; striosomes −5.4 ± 0.6 mV, n=56; matrix, −1.8 ± 0.8 mV, n=15; p=0.0114, Wilcoxon Rank Test) (Figures 1G and 1H). Therefore, our data show that relative to the matrix compartment, the striosome compartment represents the strongest source of inhibition from dorsal striatum onto the SNc dopamine neurons.

The globus pallidus external segment (GPe) also has multiple genetically defined subpopulations (Hernández et al., 2015; Mallet et al., 2012; Mastro et al., 2014). To characterize inhibition from GPe subpopulations onto SNc dopamine neurons, we injected AAV1-FLEX-hSyn-CoChR-GFP into the GPe of parvalbumin (PV)-Cre and Lhx6-Cre mice (Mastro et al., 2014). We observed that the PV-positive axons filled the SNr, while the Lhx6-positive axons invaded the SNc layer more thoroughly (Figure 1E). In contrast to a pervious functional study in rats (Oh et al., 2016), we found that activation of PV-positive GPe axons only weakly inhibited SNc neurons. However, optogenetic activation of Lhx6-positive axons resulted in strong inhibition of tonic firing and more effective hyperpolarization of SNc neurons than activation of PV-positive axons (normalized spike frequency during inhibition as percent baseline; PV 84.0 ± 6.89% n=24, Lhx6, 38.2 ± 7.47% n=26, p<0.0001, Wilcoxon Rank Test; avg Vm during inhibition relative to baseline; PV, −0.3 ± 0.2 mV, n=24; Lhx6, −2.8 ± 0.6 mV, n=26; p=0.0011, Wilcoxon Rank Test) (Figures 1I and IJ). Therefore, the Lhx6-positive neurons are the stronger source of GPe inhibition onto SNc dopamine neurons.

To better understand the underlying differences in inhibitory efficacy between genetically-defined neural populations, we examined the short-term plasticity of synapses by testing light-activated synaptic currents in SNc neurons. We found that the striosome and matrix axons both make functional synapses onto the SNc dopamine neurons, generating inhibitory post-synaptic currents (IPSCs). The currents evoked by the optical stimulus exhibited a fast, transient component and a slow, tonic component that increased in amplitude throughout the stimulus train. Stimulation of striosomal axons showed facilitation of the transient IPSCs, while stimulation of matrix axons showed no short-term plasticity (Figure 1M). We observed the slow tonic current only following stimulation of striosomal axons, but not matrix axons (Figure 1N). These results demonstrate that there are fundamental differences between striosomal and matrix synaptic connections with dopamine neurons.

Inhibitory currents from both GPe populations strongly depressed and had no slow tonic current (Figures 1L – 1N). Activation of PV axons only resulted in IPSCs in 42% (11/26) of recorded SNc neurons, while activation of Lhx6 axons showed 100% connectivity (28/28 neurons). In addition, the amplitude of the first IPSC from PV axons was significantly smaller than that of Lhx6 axons (avg amplitude PV: 52.5 ± 9.95 pA, n=11; Lhx6: 116 ± 17 pA, n=28, p=0.024, Wilcoxon rank test). Therefore, the difference in inhibitory efficacy between the GPe populations is primarily due to a higher level of connectivity from Lhx6-positive neurons onto SNc neurons, consistent with the axonal projection pattern (Figure 1E). These results demonstrate that the Lhx6 and PV synaptic connections to SNc dopaminergic neurons are similar in type but differ in connectivity strength.

### Striosomal input induces dopamine rebound

SNc dopamine neurons have been shown to rebound following aversive pauses in activity (Fiorillo et al., 2013b, 2013a; Lerner et al., 2015). To determine the ability of these inhibitory inputs to evoke rebound activity, we measured the instantaneous action potential frequency during tonic firing before optogenetic activation of either striosomal or GPe axons (PV and Lhx6 were pooled) and compared it to the instantaneous frequency of the first interspike interval after release of inhibition. In the subset of cells that were successfully inhibited, we found that inhibition from striosomal axons resulted in significantly higher rebound frequencies than inhibition from GPe axons (avg rebound frequency, striosomes: 5.31 ± 0.40 Hz, n=36; GPe: 2.75 ± 0.3 Hz, n=15; p=0.0004, Wilcoxon Rank Test, Figures 2B – 2C). Therefore, inhibition from striosomes, but not GPe, induces rebound firing in SNc dopamine neurons.

**Figure 2.**
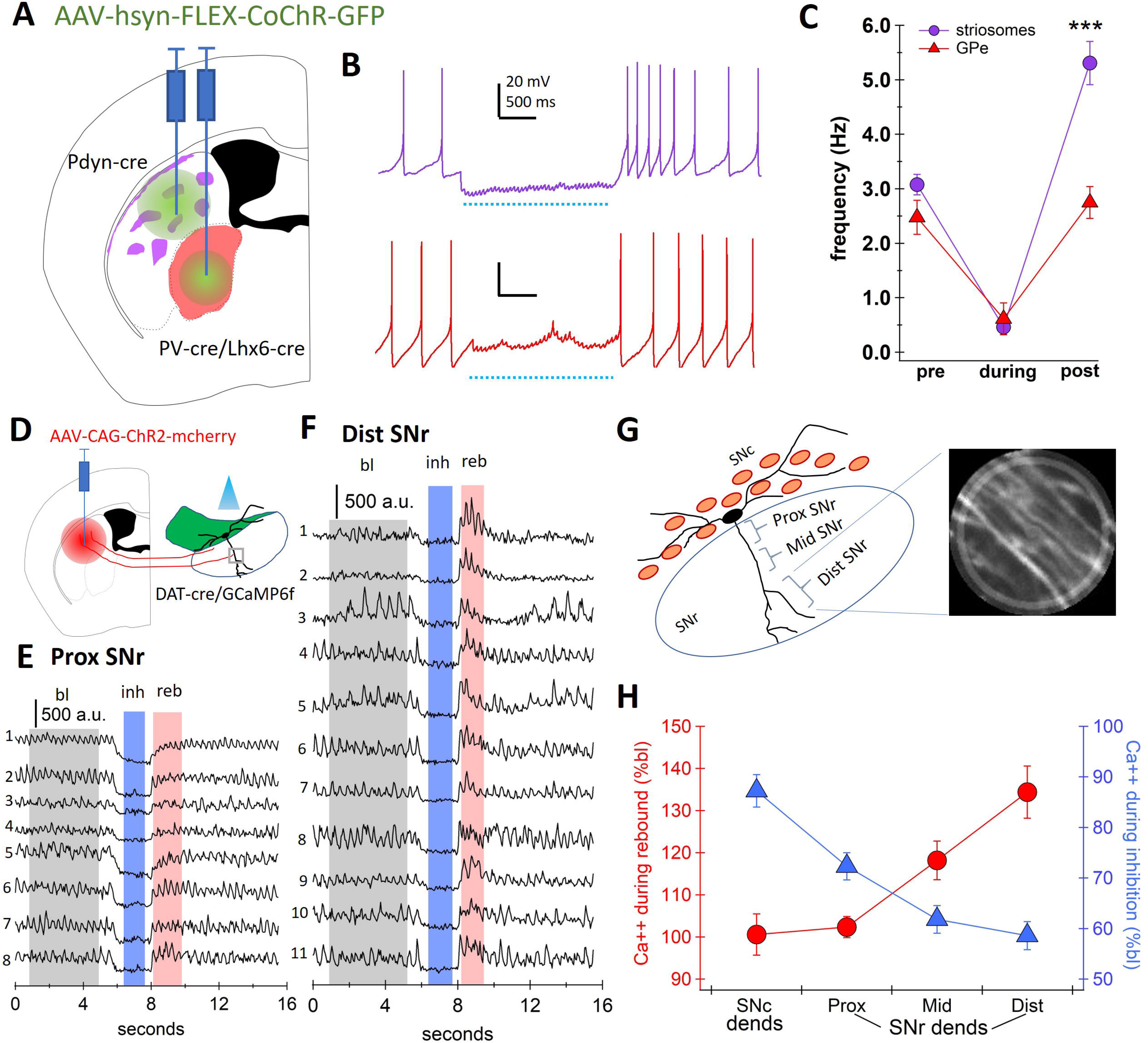
Striosomal input induces dopamine neuron rebound. **A.** Schematic of injection locations showing striatal patches (striosomes) and globus pallidus (GPe). **B.** Example current clamp traces from SNc dopamine neurons, where spontaneous tonic firing is inhibited by optogenetic activation of axons from striosomes (top, purple) or globus pallidus (bottom, red). Note rebound increase in firing frequency after striosomal input, but not GPe input. **C.** Average frequency before (pre), during, and immediately after (post) optogenetic activation of inhibitory axons. **D.** Schematic of striatal injection site in striatum of DAT-Cre/GCaMP6 mice. Note that in D-H, dorsal striatum projections from both striosome and matrix are ChR2 positive. **E.** GCaMP6f calcium signals from 8 individual reticulata-located ‘SNr dendrites’ imaged within the same region of interest (ROI). ROI located in the proximal (0-100 microns from SNc cell body layer) substantia nigra pars reticulata (SNr). Baseline shaded gray, inhibition period shaded blue, rebound period shaded red. **F.** Same as E, but for 11 individual SNr dendrites imaged in the distal (>200 microns from SNc cell body layer) region of the pars reticulata. **G.** Schematic of SNc dopamine neuron with laterally-projecting ‘SNc dendrites’ and a ventrally-projecting ‘SNr dendrite’ (left). Two-photon image of dopamine neuron dendrites located in the SNr (right). **H.** Average calcium signal as percent baseline during optogenetic activation of striatal axons (blue) and immediately after (red) for SNc dopamine neuron dendrites located in the SNc (SNc dends), proximal SNr (0-100 microns, Prox), middle SNr (100-200 microns, Mid), and distal SNr (>200 microns, Dist). ***p<0.001

SNc dopamine neurons have dendrites projecting ventrally into the SNr (“SNr dendrites”) and dendrites projecting along the SNc cell body layer (“SNc dendrites”) which have distinct dendritic morphologies that may influence their ability to integrate synaptic input (supplemental Figure S1). To investigate the relative contributions of these compartments to dopamine neuron rebound activity, we measured calcium activity in the SNc and SNr dendrites using DAT-Cre/GCaMP6f mice. A non-Cre dependent AAV-ChR2-mCherry was injected into the striatum. Striatal axons were activated by a blue (473 nm) laser at 20 Hz for 2 seconds. Calcium oscillations were recorded using two-photon (980 nm) scanning microscopy (Figures 2E – 2G). In distal dendrites, we observed clear inhibition in response to optical stimulation and synchronous calcium rebound when inhibition was released (Figure 2F, supplemental movie 1). Dendrites were classified as being within the SNc cell body layer (SNc dends, n=23), between 0-100 microns from the cell body layer in the SNr (prox SNr, n=69), 100-200 microns (mid SNr, n=76), and >200 microns (dist SNr, n=66) (Figures 2E – 2F). We found that both inhibition of the calcium signal and strength of the calcium rebound increased with distance from the SNc cell body layer (Figures 2G – 2H). These results suggest that intrinsic rebound mechanisms are most strongly recruited on the distal SNr dendrites.

### Striosomes functionally inhibit the ventrally-projecting SNr dendrites of dopamine neurons

Because striatal inhibition of the dendritic calcium signal was most effective on the distal SNr dendrite, we tested whether the density of striosomal synapses differed with dendrite type and with distance from the soma. To map striosomal synaptic location, we injected Cre-dependent synaptophysin-mCherry into the striatum of pdyn-Cre mice. In slices from these mice, we filled dopamine neurons with neurobiotin. Using tissue clearing and confocal imaging, we imaged the dopamine neuron with the mCherry-synaptophysin puncta, reconstructed each dopamine neuron (n=10), and manually identified points of overlap between puncta and dendrites at each location along the dendrite (Figures 3B – 3C). Using Sholl analysis for quantification, we found that synaptic density increased with distance from the soma along the SNr dendrite (proximal (at 50μm from soma) 2.0 ± 0.69 puncta per 10μm; distal (at 250μm from soma) 3.8 ± 0.75 puncta per 10μm), but synaptic density was uniformly low along the SNc dendrite (proximal (at 50μm from soma) 0.49 ± 0.18 puncta per 10μm; distal (at 250μm from soma) 0.23 ± 0.24 puncta per 10μm) (Figure 3C). These findings show that the ventrally-projecting SNr dendrites of SNc dopamine neurons receive dense synaptic input from striosomes.

**Figure 3.**
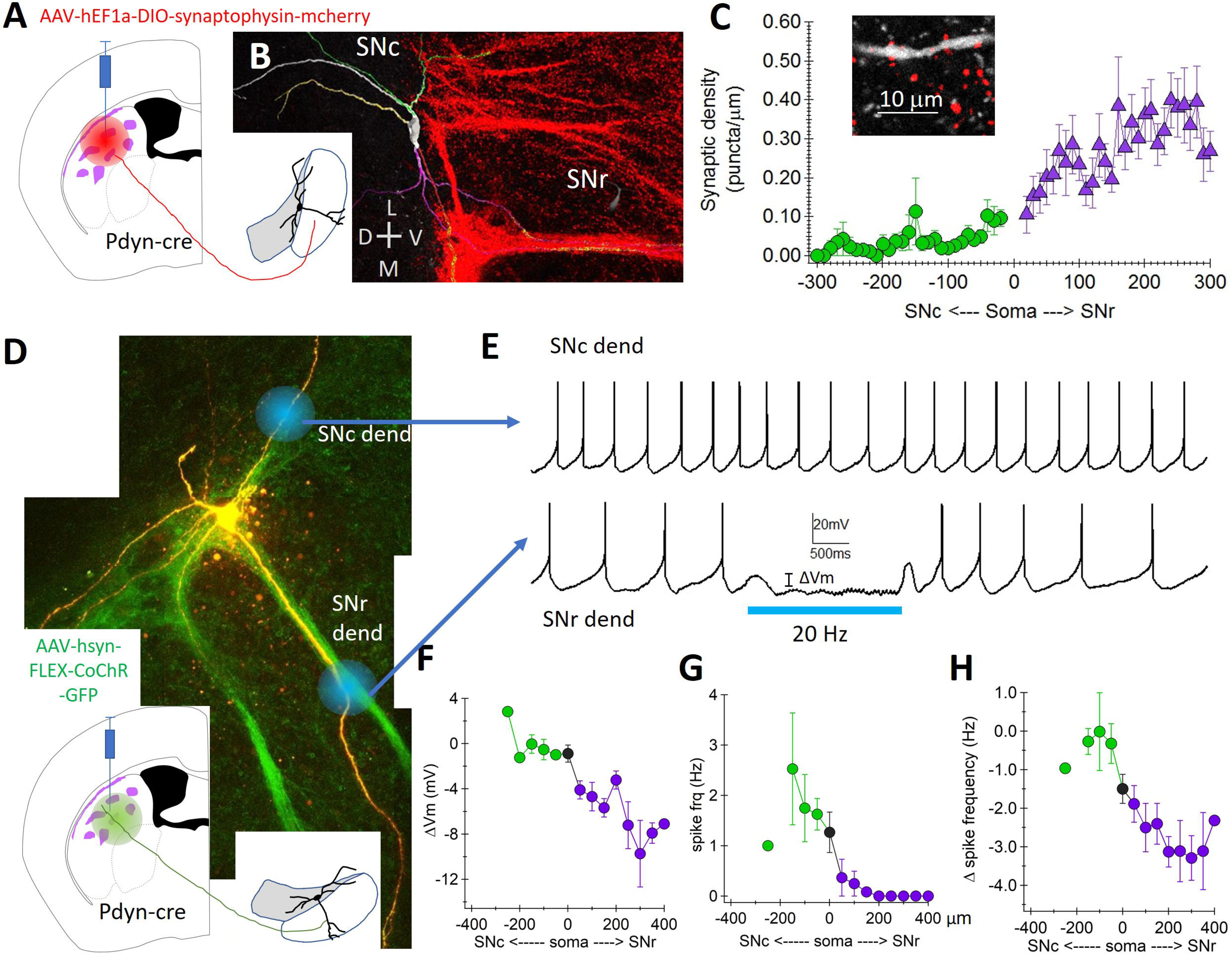
Striosomal projections selectively inhibit ventral SNr-located dendrites of SNc dopamine neurons. **A.** Schematic of injection site in striatum. **B.** Compressed Z-stack image of mCherry-labeled synaptophysin puncta (red) from striosomes, along with image of reconstructed dopaminerigc neuron with SNr dendrite. **C.** Puncta density quantified for the ‘SNc-located dendrites’ (green) and the ‘SNr-located dendrites’ (purple). Inset: single plane image of neurobiotin-filled dopamine neuron dendrite (white) and synaptophysin puncta from striosomes (red). **D.** Two-photon image of filled SNc dopamine neuron (yellow) with striosomal axons (green). Blue dots indicate locations of one-photon spatially-specific optogenetic activation of striosomal fibers. Inset: schematic of injection site. **E.** Example traces of dopamine neuron activity in response to local optogenetic activation of striosomal axons at ‘SNr dendrite’ location (top) and ‘SNc dendrite’ location (bottom). Traces recorded from neuron shown in D. **F.** Summary of membrane potential hyperpolarization (∆Vm) with distance (microns) from the soma along SNr and SNc dendrites. **G.** Action potential frequency during optical activation of striosomal axons on SNr and SNc dendrites. **H.** Change in action potential frequency from baseline firing rate during optical activation of striosomal fibers along SNr and SNc dendrites.

To examine the efficacy of inhibition as a function of dendritic location, we used spatially-specific, one-photon laser activation (473nm) to stimulate striosome fibers along SNc and SNr dendrites of a single dopamine neuron recorded in current-clamp (Figure 3D). We found that laser spot activation of striosomal axons on the distal (>150 μm from the soma) SNr dendrite completely stopped firing in all cells (7/7), while the laser spot activation on the SNc dendrite was ineffective (Figures 3E-3H). In addition, the magnitude of the somatic hyperpolarization increased with distance from the soma along the SNr dendrite (Figures 3E – 3F). These results show that the striosome axons strongly and selectively inhibit the ventrally-projecting SNr dendrite of SNc dopamine neurons.

### Striosomes activate GABA-A and GABA-B receptors on the SNr dendrite

To determine which receptor types were activated at each location along the dendrites, we measured the currents in response to activation of striosomal axons (5 pulses, 20 Hz). To ensure spatial specificity, we applied TTX (0.5 μM) and 4-AP (300 μM) to block voltage-gated sodium and potassium channels respectively, preventing action potential propagation along the axons. Voltage clamping SNc neurons, we measured IPSCs in response to laser (5 pulses 20 Hz) stimulation at multiple locations along the SNc and SNr dendrites (Figure 4A). We found that the initial fast IPSC is larger on the SNr dendrite than on the SNc dendrite (Figure 4B). To determine which synaptic receptors contributed to the IPSCs activated by striosome projections, we applied gabazine (GZ, 10 μM) to selectively block GABA-A receptors. In the presence of GZ, transient IPSCs were eliminated (average amplitude of transient current: Control 77.1 ± 24.8 pA, GZ 9.77 ± 0.971 pA, p=0.0078 Wilcoxon Signed Rank test, n=9, paired), while the slow tonic current persisted. This tonic current was eliminated by CGP55845 (1 μM), a GABA-B antagonist (average amplitude of tonic current: Control 62.6 ± 18.9 pA, GZ 25.7 ± 7.09 pA, GZ+CGP 0.79 ± 0.848 pA, control vs GZ p=0.0039, GZ vs GZ+CGP p=0.0039, Wilcoxon Signed Rank test, n=9 paired) (Figure 4C). In the presence of GZ, the amplitude of the isolated GABA-B IPSC increased with distance from the soma on the SNr dendrite (Figure 4D).

**Figure 4.**
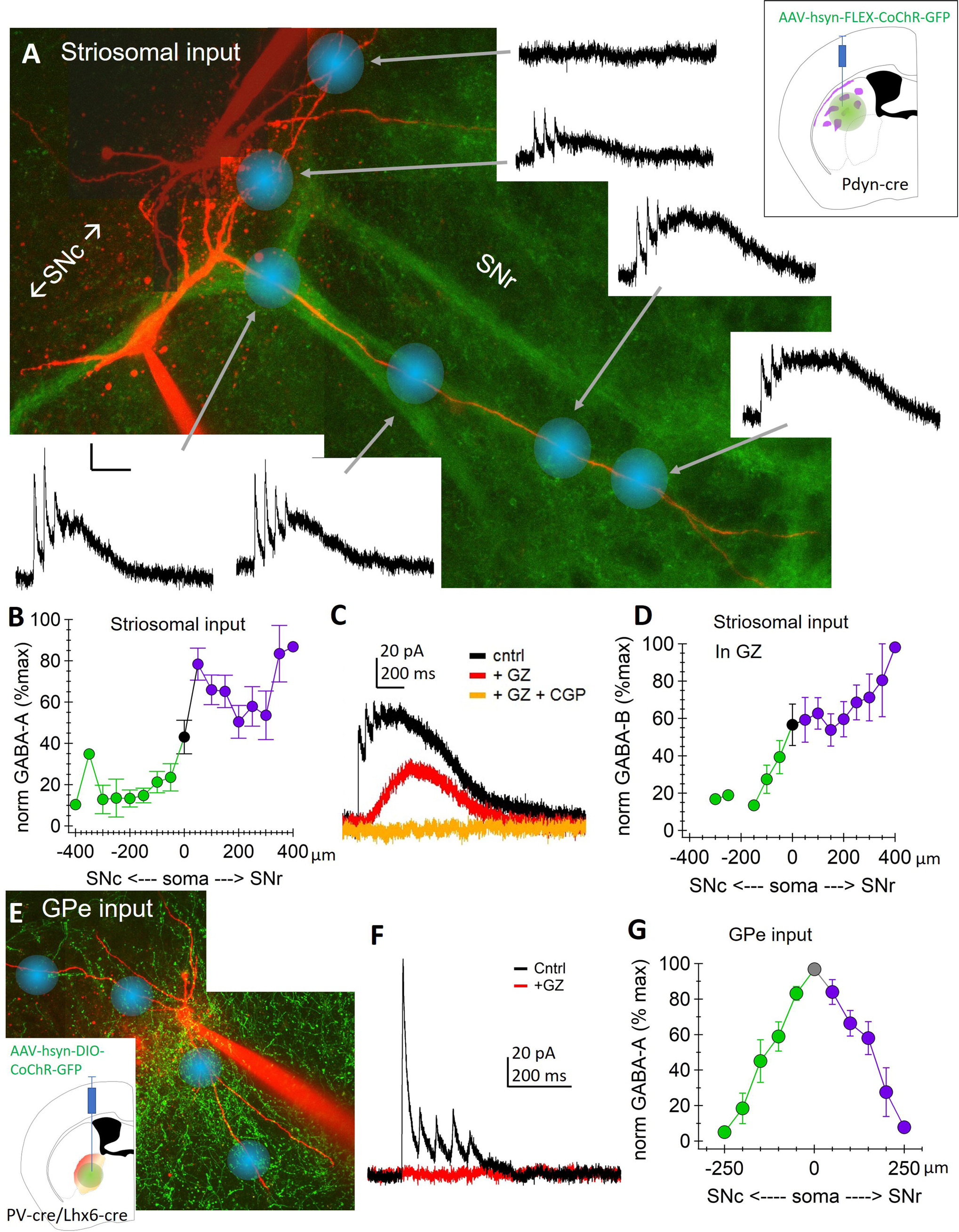
Striosomal input activates GABA-A and GABA-B receptors on the SNr-located dendrite, while GPe input activates GABA-A on the soma and proximal dendrites. **A.** Two-photon image of Alexa-594 filled SNc dopamine neuron (red) and striosomal axons (green). Blue spots indicate locations of focal optogenetic activation of striosomal axons (5 light pulses, 20Hz, in the presence of 0.5 μM TTX and 300 μM 4-AP). The corresponding inhibitory current traces are connected to each location with gray arrows. Second neuron in image was not successfully recorded and has been darkened for clarity. Inset: schematic of injection site in striatum. Scale bars: 20pA, 200ms. **B.** Summary of normalized transient current for the first stimulus of train. Currents plotted against distance from soma (in microns) along the SNr dendrite (right) and the SNc dendrite (left) and normalized to maximal current amplitude for each cell. Note, averaged maximal current recorded when light spot placed on SNr dendrite. **C.** Example traces showing pharmacological effect of 10 μM gabazine (GZ) to block fast GABA-A mediated currents and 1 μM CGP55845 (CGP) to block slow GABA-B mediated tonic current evoked by optogenetic activation of striosomal axons. **D.** Same as B, showing the normalized peak amplitude of the isolated GABA-B current (recorded in GZ) normalized to the amplitude of the maximum current for each cell. **E.** Two-photon image of SNc dopamine neuron (red) and globus pallidus (GPe) axons (green). Inset: schematic of injection site in GPe. **F.** Example traces showing synaptic currents evoked by optogenetic activation of GPe inputs in control (black) and in the presence of gabazine (red). **G.** Summary of normalized transient current amplitude by distance from the soma (in microns) along the SNc (left) and SNr (right) dendrites. Currents in B, D, and G are normalized to maximal current amplitude for each cell.

For comparison, we tested the functional location of GPe inputs using this same method. For both PV and Lhx6 inputs, we observed fast transient currents (average amplitude of transient current: 114 ± 17.6 pA) with little to no slow tonic current (average amplitude of tonic current: 7.87 ± 2.55 pA) (Figure 4F). Application of GZ abolished the entire synaptic current, consistent with GPe inputs activating mainly GABA-A receptors. The transient component was consistently large at the soma and became progressively smaller in amplitude along both the SNr and SNc dendrites. Because there was no location difference between the PV (n=5) and Lhx6 (n=6) inputs, they were pooled (Figure 4G). Together, these experiments show that inputs from striosomes onto SNc dopamine neurons activate both GABA-A and GABA-B receptors on the ventrally-projecting SNr dendrites, while the inputs from the GPe selectively inhibit the soma and proximal dendrites.

### Computational modeling shows that striosomal synaptic characteristics are optimized to induce rebound

To examine the synaptic characteristics of the striosomal input that contribute to rebound, we generated a multi-compartmental computational model of an SNc dopamine neuron based on our neural reconstructions (Supplemental Figure S1). Striosomal synapses were simulated by combining a facilitating GABA-A conductance with a slow GABA-B conductance (Beckstead and Williams, 2007) (Figure 5B, above). GPe synapses were simulated as a GABA-A only conductance with synaptic depression (Figure 5C, below). Striosomal synapses were placed on the SNr dendrites and GPe synapses on the soma and proximal dendrites (Figure 5A). To explore the effect of hyperpolarization on rebound, we generated an inhibition-rebound curve by adjusting the maximal inhibitory conductance. Our simulations showed a strong relationship between hyperpolarization and rebound for striosomal inhibition, while GPe inhibition did not generate rebound, consistent with our experimental findings (Figures 5C and 5D). This occurs because hyperpolarization from GPe inputs is limited by the depolarized reversal potential of GABA-A receptors (Erev = −65 mV). By contrast, activation of striosome inputs involves GABA-B receptors which more strongly hyperpolarize through inwardly rectifying G-protein coupled potassium channels (GIRKs), allowing for recruitment of T-type calcium channels and hyperpolarization-activated cation currents (Ih) current (Evans et al., 2017). These simulations show that the striosomal synaptic characteristics are sufficient to generate rebound in a model SNc dopamine neuron.

**Figure 5.**
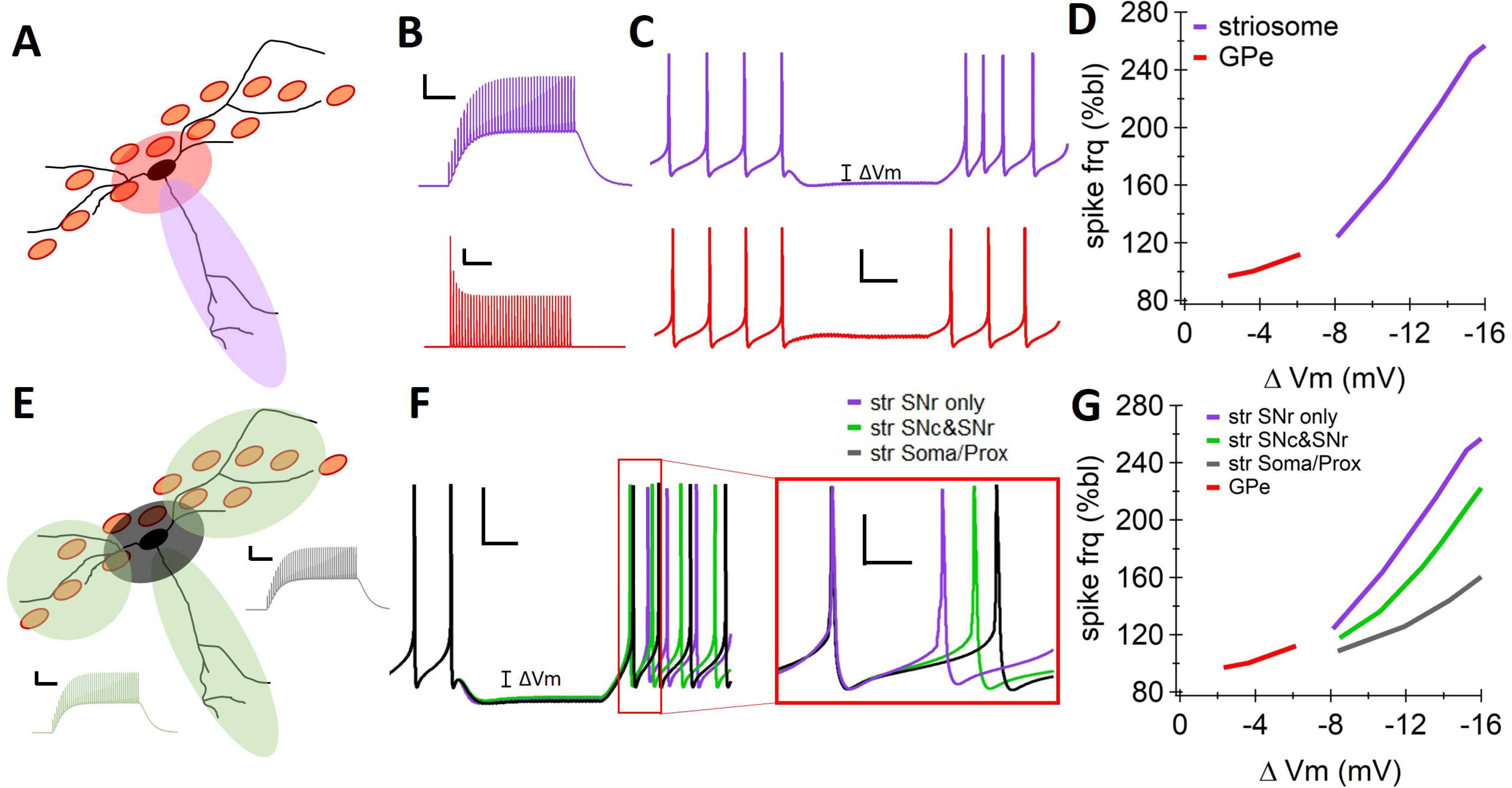
Computational modeling shows that striosomal input is synaptically optimized to induce rebound. **A.** Schematic of model SNc dopamine neuron with SNc- and SNr-located dendrites. Shaded regions indicate location of simulated synaptic inhibitory input from the GPe (red) and the striosomes (purple). **B.** Simulated synaptic conductances for a 20 Hz, 2 second train for striosomal (top, scale bar 50 pS, 500 ms) and GPe (bottom, scale bar 500 pS, 500 ms). **C.** Simulations of dopaminergic neuron response to striosomal input (top) and GPe input (bottom). Scale bars 20 mV, 500 ms. Note rebound increase in action potential frequency from striosomal, but not GPe input. **D.** Graph of normalized frequency during rebound (normalized to baseline firing rate) plotted against membrane hyperpolarization in response to striosome (purple) or GPe (red) input. **E.** Schematic showing dendritic and somatic locations of striosomal input for simulations in F and G. Insets show the same striosomal characteristics (as in C) were used. **F.** Example simulations from striosomal input on SNr dendrite only (purple, as in C), on all dendrites (green), and on the soma and proximal dendrites (black). Scale bars 20 mV, 500 ms. Inset: closeup of rebound firing, traces aligned to first action potential, scale bars: 20 mV, 100 ms. **G.** Graph of normalized rebound firing plotted against somatic hyperpolarization for different arrangements of striosomal input and GPe input. Note, relationship is steepest when striosomal input is located on the SNr dendrites only.

To determine how dendritic location of inhibitory input contributes to rebound firing, we placed striosomal inputs in three spatial configurations – 1) SNr dendrite only, 2) perisomatic or 3) all dendrites (Figure 5E). When measured from the same somatic potential, striosomal inhibition of the SNr dendrites alone induced the largest increase in rebound frequency, while perisomatic inhibition induced the weakest (Figure 5F). The relationship between somatic hyperpolarization and rebound frequency was steepest when the SNr dendrites were selectively inhibited, and shallow when striosomal inhibition was located on the soma and proximal dendrites (Figure 5G). Therefore, these simulations show that the dendrite-specific nature of the striosomo-nigral synapses amplifies their ability to induce rebound firing in SNc dopamine neurons.

### Striosomes selectively inhibit the ‘rebound-ready’ subset of SNc dopamine neurons

The SNc can be divided into a ventral tier that is positive for Aldh1a1 and a dorsal tier which is positive for calbindin (Gerfen et al., 1987; Kim et al., 2015; Poulin et al., 2018; Wu et al., 2019). We have previously shown that the ventral tier SNc neurons rebound more strongly from hyperpolarization due to their strong expression of T-type calcium channels and large Ih current (Evans et al., 2017). To compare the strength of striatal inhibition onto dorsal and ventral tier SNc neurons, we optically stimulated striatal inputs and imaged somatic calcium signals in SNc neurons from DAT-Cre/GCaMP6f mice (Figure 6A). We found that striatal inhibition effectively reduced calcium signals in ventral tier neurons but only weakly inhibited calcium signals in dorsal tier neurons (n=446 cells from 5 slices from 3 mice; Figures 6B – 6D). Therefore, these results demonstrate that striatal inhibition is limited to a subpopulation of ventral tier SNc neurons (Figure 6D).

**Figure 6.**
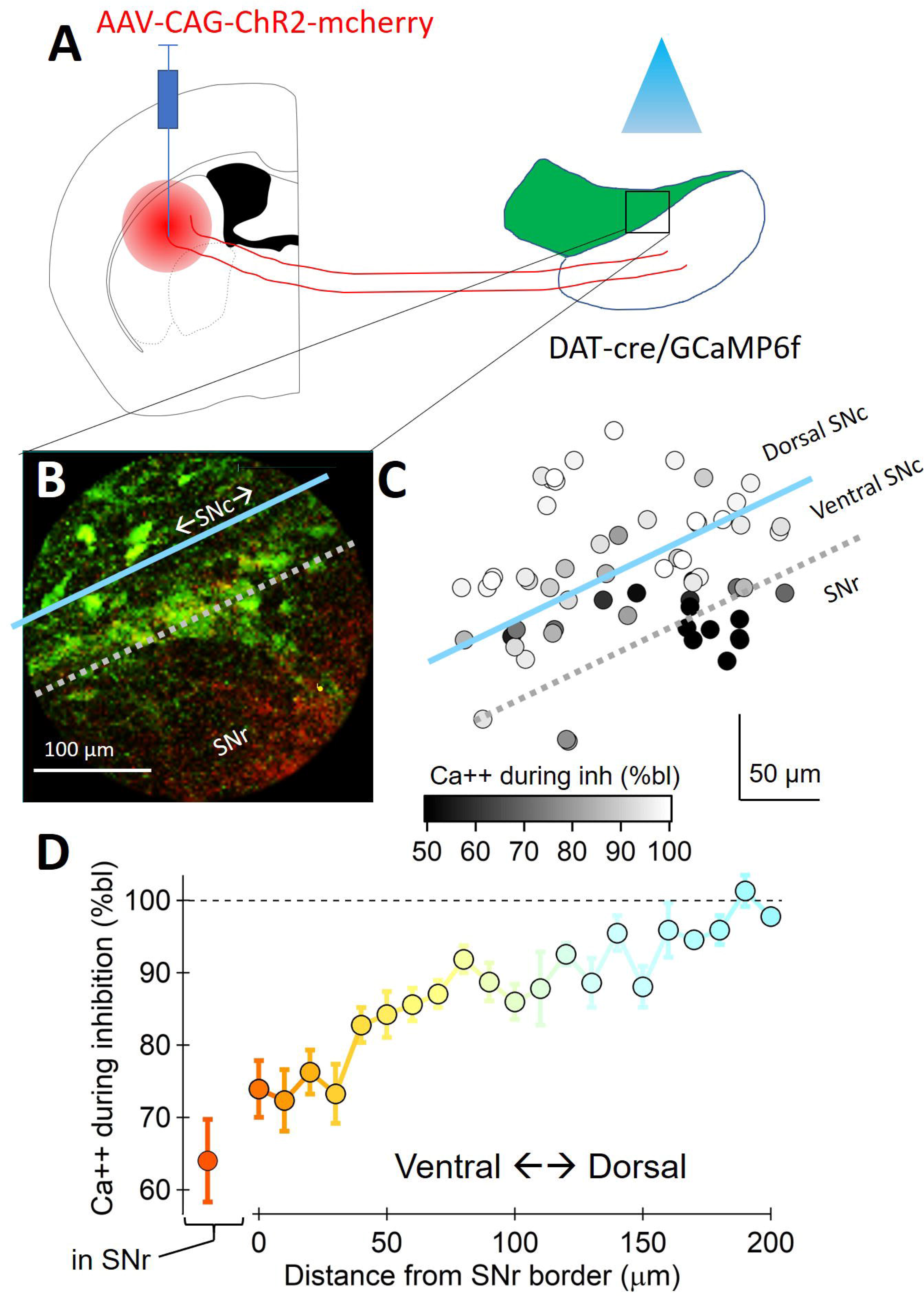
Striosomes selectively target a subset of ventral SNc dopamine neurons. **A.** Schematic of striatal injection site in dorsal striatum of DAT-Cre/GCaMP6f mouse. **B.** Two-photon image of GCaMP6f-positive SNc dopamine neurons (green) with ChR2-mCherry infected striatal axons (red). **C.** SNc dopamine neuron cell bodies color coded by calcium reduction (% baseline, bl) due to optogenetic activation of striatal axons. In B and C, blue line divides dorsal and ventral SNc, and dotted gray line indicates the border between the SNc cell body layer and the SNr. **D.** Amount of calcium inhibition (% baseline) due to optogenetic activation of striatal axons with distance from the SNc-SNr border. Ventral neurons are more strongly inhibited than dorsal neurons.

To further understand the subpopulation targeted by striosomes, we explored the relationship between striosomal inhibition and dendritic configuration. We found that striosomal input strongly inhibited some SNc cells but only weakly inhibited others. To account for these differences, we separated dopamine neurons into three morphologically-defined groups: neurons with dendrites in striosome-dendron bouquets (Morph 1; n=36), SNr dendrites that do not participate in bouquets (Morph 2; n=13) or no SNr dendrite (Morph 3; n=5) (Figures 7A – 7B). SNc neurons that had somas located at the tops of bouquets but did not have clear dendrites in bouquets (4 out 36 cells) were considered bouquet-participating cells. We found that dopamine neurons participating in bouquets have reduced spiking during inhibition and are more strongly hyperpolarized than neurons in the other two groups (avg spike frequency during inhibition, Morph 1: 0.56 ± 0.17 Hz, Morph 2: 1.8 ± 0.45 Hz, Morph 3: 2.5 ± 0.49 Hz; p=0.007 Kruskal-Wallis test; Morph 1 vs Morph 2: p=0.04; Morph 1 vs Morph 3: p=0.001; Wilcoxon rank test; avg Vm hyperpolarization, Morph 1: −7.1 ± 0.8 mV, Morph 2: −3.5 ± 1.3, Morph 3: −0.2 ± 0.1 mV; p=0.0007 Kruskal-Wallis test; Morph 1 vs Morph 2: p=0.023; Morph 1 vs Morph 3: p=0.00011; Wilcoxon Rank Test) The rebound spike frequency was also highest in bouquet-participating neurons (Avg spike frequency during rebound: Morph 1: 4.97 ± 0.44 Hz, Morph 2: 3.56 ± 0.43 Hz, Morph 3: 2.46 ± 0.57 Hz; p=0.02, Kruskal-Wallis test. Morph 1 vs Morph 2 p=0.078; Morph 1 vs Morph 3 p=0.014, Wilcoxon Rank Test) (Figures 7C–7D). These results show that striosome-dendron bouquets are sites of particularly strong striatonigral inhibition.

**Figure 7.**
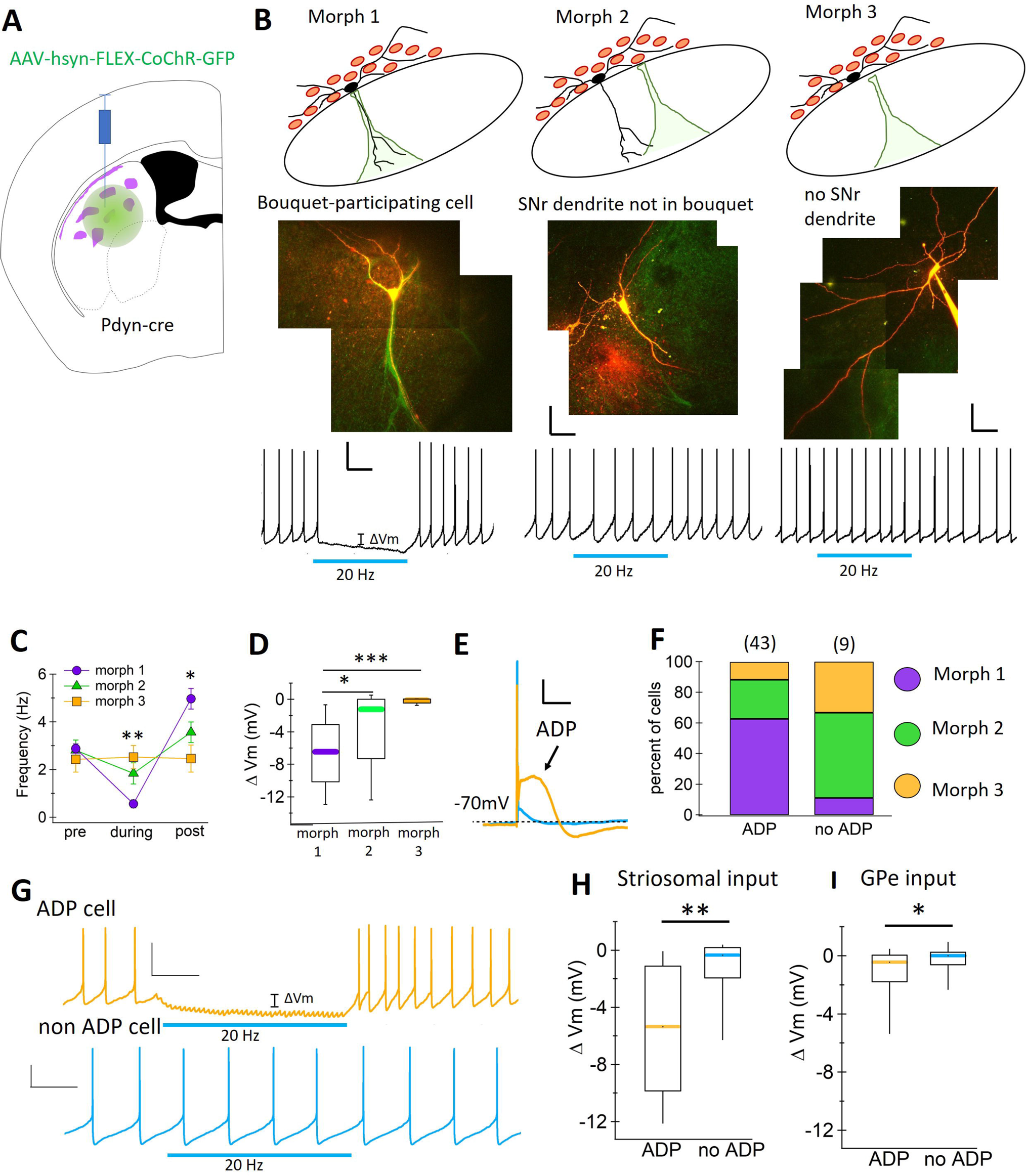
Striosomes preferentially inhibit the ‘rebound-ready’ SNc neurons. **A.** Schematic of injection site in dorsal striatum. **B.** SNc dopamine neurons divided into three morphological categories: Morph 1, neurons with a dendrite, soma or both located within a striosome-dendron bouquet; Morph 2, neurons with a dendrite in the SNr, but not participating in a bouquet; Morph 3, cells with no dendrite in the SNr and soma not located in a bouquet. Top: schematic of each morphology type, green shading represents striosomal innervation of dendron bouquet. Middle: Example two-photon image of each cell type (red) with striosomal axons (green). Bottom: Example traces from pictured cells showing response to optogenetic activation of striosomal axons during tonic firing. Scale bar 20 mV, 500 ms. **C.** Summary of action potential frequency before (pre), during (dur), and after (post) optogenetic activation of striosomal axons for each morphology type. **D.** Membrane potential hyperpolarization during optogenetic activation of striosomal axons compared to baseline for each morphology. **E.** Example traces from one SNc dopamine neuron with a low-threshold afterdepolarizations (ADP), and one without (non-ADP). Scale bars 20 mV, 100 ms. **F.** Dopamine neurons with ADPs are more likely to be Morph 1 neurons than those without ADPs. **G.** Same neurons as in E, showing response to optogenetic activation of striosomal axons. **H.** ADP cells are more strongly inhibited by optogenetic activation of striosomal axons than non-ADP cells. **I.** ADP cells are only slightly more hyperpolarized by globus pallidus (GPe) input than non-ADP cells. *p<005, **p<0.01, ***p<0.001.

Finally, we directly classified SNc neurons based on their intrinsic rebound mechanisms. An electrophysiological signature of the ‘rebound-ready’ SNc neurons is that they exhibit an after-depolarization (ADP) in response to stimulation at hyperpolarized potentials (Evans et al., 2017). To test whether the striosomal input preferentially inhibits the intrinsically ‘rebound-ready’ SNc neurons, we separated SNc neurons into ADP cells (rebound-ready) and non-ADP cells (non-rebounding) (Figure 7E). Analyzing the morphology of each group, we found that ADP cells were more likely to have dendrites in bouquets than non-ADP cells (percent bouquet-participating cells, ADP: 62.8%, 27/43 cells; non-ADP: 11.1%, 1/9 cells) (Figure 7F). Similarly, we found that striosomal inhibition was stronger on ADP cells than on non-ADP cells (avg Vm vs baseline, ADP: −5.9 ± 0.74 mV, n=45; non-ADP: −1.2 ± 0.78 mV, n=8; p=0.0089, Wilcoxon rank test) (Figures 7G-7H). By contrast, we found that GPe inhibition was only slightly stronger onto ADP cells compared to non-ADP cells (avg Vm vs baseline, ADP: −2.2 ± 0.54 mV, n=33; non-ADP: −0.4 ± 0.31 mV, n=17; p=0.017, Wilcoxon Rank test) (Figure 7I). Together, these results show that striosomes preferentially inhibit the rebound-ready subset of SNc dopamine neurons.

## Discussion

In this study, we have defined a distinct striatonigral circuit which facilitates dopamine neuron rebound activity. Specifically, the striosomes of the dorsal striatum preferentially inhibit the ventral tier SNc dopamine neurons which exhibit strong intrinsic rebound properties. This striosomal input is located on the SNr dendrite and activates both GABA-A and GABA-B receptors. Our computational modeling indicates that the dendrite-specific location of the striosomal input contributes to its ability to generate rebound. Importantly, striosome-induced rebound activity likely involves the interplay of GABA-B receptor activated potassium current (GIRK) which hyperpolarizes cells to recruit hyperpolarization-activated cation currents (Ih) and T-type calcium currents that trigger rebounds (Evans et al., 2017). By contrast, inputs from the GPe onto SNc dopamine neurons do not activate GABA-B receptors, are not dendrite specific, and do not induce rebound (Figure 8). Therefore, the striosomo-nigral-striatal connection represents a self-contained circuit mechanism by which striatal neurons can trigger rebound-induced phasic increases in striatal dopamine without the need for external excitatory input.

**Figure 8.**
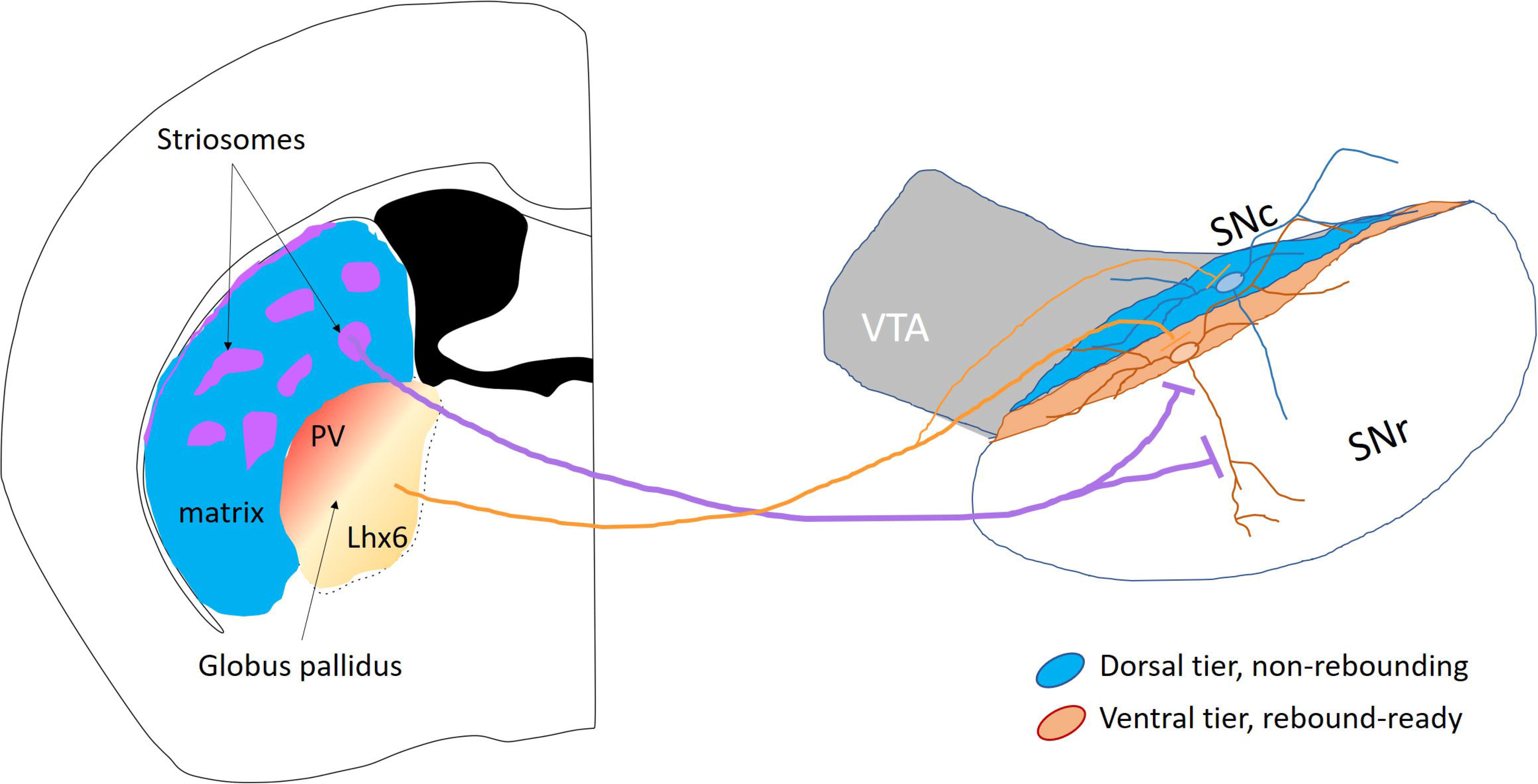
A distinct striosomo-nigral circuit facilitates dopamine rebound. Striosomal input to SNc dopamine neurons is synaptically optimized to induce rebound and preferentially inhibits the SNc dopamine neurons that are intrinsically able to rebound.

### Function of genetically-defined inhibitory inputs onto SNc neurons

Previous *in vivo* work has shown complex multi-phasic responses in SNc dopamine neurons upon electrical stimulation of the striatum or GPe (Brazhnik et al., 2008; Paladini et al., 1999). By testing inputs onto SNc neurons from genetically-defined inputs in isolation, we show that striosomal inputs are the predominate source of inhibition from the striatum and Lhx6-positive inputs are the predominate source of inhibition from the GPe. Although both inputs could pause firing, striosomal currents were dendrite-specific and showed synaptic facilitation while GPe currents were somatic and showed synaptic depression. These short-term plasticity results are similar to previous findings comparing striatal and pallidal inputs to SNr GABAergic neurons (Connelly et al., 2010). These opposite spatial and temporal characteristics suggest that GPe input is optimized to communicate fast, sudden signals to pause dopamine neurons only transiently, while striosome input is optimized to communicate more sustained signals.

Experiments in behaving animals show that striatal projection neurons fire bursts of action potentials up to 16 Hz *in vivo* (Sippy et al., 2015). However, the strength of the striatal inhibition onto SNc dopamine neurons is likely not due to the firing frequency of one cell, but rather may rely on high levels of convergence (Watabe-Uchida et al., 2012). Currently, it is unclear how many striosomal neurons participate in a single bouquet structure, or whether a bouquet is formed by a single striosome or many. Axon tracing in monkeys shows that multiple neurons from a single striosome send axons into several distinct bouquet-like structures (Lévesque and Parent, 2005). Because there are several types of striosomes (Davis et al., 2018; Miyamoto et al., 2018), it will be important to determine whether striosomes act as isolated units (Banghart et al., 2015) or communicate cohesive signals.

An important function of striosomes may be to synchronize the activity of SNc dopamine neurons. Our calcium imaging shows that dendrites within bouquets exhibit spontaneous asynchronous calcium oscillations under baseline conditions but show synchronous increases in calcium when striatal inhibition is released (supplemental movie 1). The strong rebound characteristics of the striosome-inhibited subpopulation of neurons will contribute to a synchronous rebound by allowing fast return to firing after inhibition. However, this characteristic is not ubiquitous among dopamine neurons, as dopamine neurons in the VTA are more variable in their recovery time from inhibition due to the slow A-type potassium current kinetics (Tarfa et al., 2017). Mechanistically, the structure of the striosome-dendron bouquet may contribute to synchrony among dopamine neurons. Single striatal axons will likely form synapses onto multiple dendrites within the bouquet, which may synchronize activity. Such a tightly packed structure would facilitate the propagation of a unified signal to all bouquet-participating cells. Therefore, the synaptic characteristics of the striosomal input and the intrinsic characteristics of the ventral SNc neurons are optimized to produce synchronous rebound activity.

### Role of GABA-B inhibition on SNc dopamine neurons

Anatomical results have shown that striatal inputs onto SNc dopamine neurons are less numerous at the soma, but a higher density on the distal dendrites (Bolam and Smith, 1990), leading to the assumption that striatal input may only weakly effect dopamine neuron activity. In contrast to this view, we demonstrate that activation of striosome projections results in strong hyperpolarizing control over the soma and prevents action potential firing in SNc neurons. The strength of the striosomal input is due to their activation of both GABA-A and GABA-B receptors. In particular, GABA-B activation strongly inhibits dopamine neurons by activating the inwardly-rectifying G-protein coupled potassium (GIRK) channels (Beckstead and Williams, 2007; Koyrakh et al., 2005) and blocking the sodium leak channel NALCN (Philippart and Khaliq, 2018). Uncaging experiments in cultured dopamine neurons show that GABA-B activation on the dendrites is more effective than GABA-A activation (Kim et al., 2018). This is likely due to the slow kinetics of the GABA-B receptor and the simple architecture the SNr dendrites (Supplemental Figure S1), which enables efficient propagation of inhibitory signals from distal dendrites to the soma.

Past work examining inhibitory inputs to the ventral tegmental area (VTA) has shown circuit specific activation of GABA-B receptors onto VTA dopaminergic neurons (Cameron and Williams, 1993; Edwards et al., 2017; Yang et al., 2018). For SNc neurons, we find that the striosome-dendron bouquets are a site of strong GABA-B receptor activation. It is unclear whether the densely packed striosomal synapses within the dendron bouquets (Figure 3; and see Crittenden et al., 2016) facilitates GABA-B receptor activation. Future work is needed to determine whether a similar organizing principle contribute to GABA-B signaling in VTA.

It has been suggested that GABA-B receptor activation would not strongly generate rebound due to its slow on and off kinetics. Our previous work showing strong rebound activity in the ventral tier SNc dopaminergic neurons used direct somatic hyperpolarization, which results in a much faster relief from inhibition than the GABA-B receptor (Evans et al., 2017). However, GABA-B antagonists infused into the SNc *in vivo* causes reduced bursting of dopamine neurons, suggesting that activation of GABA-B promotes bursting activity (Paladini and Tepper, 1999). This observation is consistent with our findings that simultaneous relief from synaptic GABA-A and GABA-B receptor-mediated inhibition generates dopamine neuron rebound. The GABA-B receptor dependent rebound activity presented here is reminiscent of the disinhibition burst firing proposed by Paladini and colleagues (Lobb et al., 2010) but differs in that it involves intrinsic rebound mechanisms and does not rely solely on synaptic input.

### Defining SNc dopamine neuron subpopulations

Past studies have classified substantia nigra dopamine neurons subpopulations according to their projection targets (Farassat et al., 2019; Lerner et al., 2015; Schiemann et al., 2012), expression of neurochemical markers (La Manno et al., 2016; Poulin et al., 2018; Wu et al., 2019), and intrinsic membrane properties (Evans et al., 2017; Neuhoff et al., 2002). Based on the anatomical results, Crittenden et al. (2016) proposed that dopamine neuron clusters (in bouquets) may form specialized nigral compartments. Our experimental findings here provide functional evidence for this hypothesis, demonstrating the presence of a novel projection-defined subpopulation of dopaminergic neurons within the SNc. We show that the influence of dorsal striatal axons is non-uniform across the SNc, preferentially innervating a subset of ventral tier SNc neurons. As described in our previous work (Evans et al., 2017), this SNc subpopulation is likely composed of calbindin-negative, Aldh1a1-positive neurons with strong intrinsic rebound properties.

Dopaminergic neurons of the SNc are also distinguished by their behavioral responses to aversive stimuli. Specifically, medial SNc neurons are inhibited by aversive stimuli while lateral neurons are activated (Lerner et al., 2015; Matsumoto and Hikosaka, 2009). However, the striosome-input defined neurons examined in this study are typically located in the middle of the SNc and therefore do not fit neatly into either medial or lateral subpopulations. These ventral SNc neurons have extensive dendrites within the SNr, which correlates with stronger inhibitory responses to aversive stimuli (Henny et al., 2012). It is also likely that this input-defined SNc subpopulation that we identify in mice is analogous the subpopulation of ventrally-located SNc neurons in monkey which rebound most strongly after an aversive event (Fiorillo et al., 2013b). Future work is needed to determine the extent to which bouquet-participating SNc neurons overlap functionally with the medial and lateral subpopulations examined in prior studies.

### What is the significance of striosomal inhibition and dopamine rebound?

Individual cells within striosomes show variable and complex responses to rewarding and aversive stimuli (Bloem et al., 2017; Yoshizawa et al., 2018). However, a subset of striosomal neurons show clear activation during an aversive air puff (Yoshizawa et al., 2018), and striosomes become over-active in conditions of chronic stress (Friedman et al., 2017). A study examining the relationship between dopamine neuron morphology and behavior found that aversive inhibition correlates with the length of the SNr dendrite and hypothesized that aversive signaling would be transmitted mainly through the SNr dendrite (Henny et al., 2012). Here, we identify the striosomes as a prominent source of inhibition onto the SNr dendrite, which reveals the possibility that they may convey an aversive signal.

Interestingly, behavioral experiments show that electrical stimulation of striosomes reinforces actions (White and Hiroi, 1998) and striosomal ablation impairs habit learning (Jenrette et al., 2019). Because dopamine is required for learning, these findings may seem counter-intuitive as we have shown that the striosomes strongly inhibit dopamine neurons. However, striosomal activation may paradoxically result in a reinforcing pulse of dopamine through disinhibition or by inducing dopamine rebound activity. Because dopamine rebound is often observed after an aversive pause in activity (Budygin et al., 2012; Fiorillo et al., 2013b; de Jong et al., 2019; Lerner et al., 2015; Wang and Tsien, 2011), rebound may represent an important learning signal signifying relief from an unpleasant stimulus and may serve to reinforce an escape behavior.

The timing of dopamine release in the striatum is a key factor in synaptic plasticity (Shindou et al., 2019; Yagishita et al., 2014). Because SNc dopamine neurons form a reciprocal loop with the dorsal striatum, dopamine rebound may represent a mechanism by which striosomes can control the timing of phasic dopamine signals in the striatum and therefore control the plasticity of their own synapses. Future work is needed to test whether self-contained dopamine rebound activity is related to the role striosomes play in repetitive behaviors (Bouchekioua et al., 2018; Canales and Graybiel, 2000) and persistent (devaluation-resistant) stimulus-response learning (Jenrette et al., 2019).

### Conclusions

We have shown that striosomes can cause a pause-rebound firing pattern in ventral dopamine neurons in the absence of excitatory input. This finding reveals a mechanism by which striosomes could control the timing of phasic dopamine signals in the striatum, potentially causing plasticity in recently activated synapses and reinforcing recent motor actions.

## Methods

### Lead contact and materials availability

Further information and requests for resources and reagents should be directed to and will be fulfilled by the Lead Contact Dr. Zayd Khaliq

### Experimental model and subject details

All animal handling and procedures were approved by the animal care and use committee (ACUC) for the National Institute of Neurological Disorders and Stroke (NINDS) at the National Institutes of Health. Mice of both sexes underwent viral injections at postnatal day 18 or older and were used for *ex vivo* electrophysiology or imaging 3-8 weeks after injection. The following strains were used: Pdyn-IRES-Cre (129S-Pdyn(tm1.1(cre)/Mjkr)/LowlJ, The Jackson Laboratory Cat#027958); Calb1-IRES2-Cre-D (129S-Calb1(tm2.1(cre)/Hze)/J, The Jackson Laboratory Cat#028532); PV-Cre (129P2-Pvalb(tm1(cre)Abr)/J, The Jackson Laboratory Cat#017320); DAT-Cre (SJL-Slc6a3(tm1.1(cre)Bkmn/J, The Jackson Laboratory Cat#006660); Ai95-RCL-GCaMP6f-D (Cg-Gt(ROSA)26Sor(tm95.1(CAG-GCaMP6f)Hze)/MwarJ, The Jackson Laboratory Cat#028865); Lhx6-Cre (CBA-Tg(Lhx6-icre)1Kess/J, obtained from the lab of Aryn Gittis).

### Method details

#### Viral injections

All stereotaxic injections were conducted on a Stoelting QSI (Cat#53311). Mice were maintained under anesthesia for the duration of the injection and allowed to recover from anesthesia on a warmed pad. The AAV-hsyn-FLEX-CoChR (Boyden, UNC vector core), AAV-CAG-hChR2-mCherry (Diesseroth, Addgene) or AAV-hEF1a-DIO-synaptophysin-mCherry (Neve, MIT Viral gene core) viruses (0.5-1μl) were injected bilaterally into either the dorsal striatum (X: ± 2.1 Y: +0.8 Z: −2.6) or the GPe (X: ± 1.9 Y: −0.5 Z: −3.9) via a Hamilton syringe. At the end of the injection, the needle was raised 1-2 mm for a 10 minute duration before needle was removed.

#### Slicing and electrophysiology

Mice were anesthetized with isoflurane and transcardially perfused with ice cold modified ACSF containing (in mM) 198 glycerol, 2.5 KCl, 1.2 NaH_2_PO_4_, 20 HEPES, 25 NaHCO_3_,10 glucose, 10 MgCl_2_, 0.5 CaCl_2_, 5 Na-ascorbate, 3 Na-pyruvate, and 2 thiourea. Mice were decapitated and brains extracted. Coronal slices were cut at 200 μm thickness on a vibratome and incubated for 30 minutes in heated (34°C) chamber with holding solution containing (in mM) 92 NaCl, 30 NaHCO_3_, 1.2 NaH_2_PO_4_, 2.5 KCl, 35 glucose, 20 HEPES, 2 MgCl_2_, 2 CaCl_2_, 5 Na-ascorbate, 3 Na-pyruvate, and 2 thiourea. Slices were then stored at room temperature and used 30 min to 6 hours later. Whole-cell recordings were made using borosilicate pipettes (2-7 MΩ) filled with internal solution containing (in mM) 122 KMeSO_3_, 9 NaCl, 1.8 MgCl_2_, 4 Mg-ATP, 0.3 Na-GTP, 14 phosphocreatine, 9 HEPES, 0.45 EGTA, 0.09 CaCl_2_, 0.05 AlexaFluor 594 hydrazide adjusted to a pH value of 7.35 with KOH. Some experiments included 0.3 mM Fluo5F in place of EGTA and CaCl_2_, and some included 0.1-0.3% neurobiotin for post-hoc visualization. Current clamp recordings were manually bridge balanced. For current clamp rebound experiments in Figure 2, inhibited cells were defined as those in which the optogenetic stimulation reduced spiking from baseline by at least 1 Hz. In voltage clamp experiments, cells were held at −50 mV and cell capacitance and access resistance (< 25 MΩ) were compensated to 30-70%. Liquid junction potential (−8 mV) was not corrected. All experiments were conducted heated (31-34°C).

#### Optogenetic activation

Experiments were conducted on an Olympus BX61W1 multiphoton upright microscope. Whole-field optogenetic activation of CoChR or ChR2 axons in brain slice was achieved by either a white LED (Prizmatix) sent through a FITC filter (HQ-FITC; U-N41001; C27045) or a blue (470nm) LED (Thorlabs, LED4D067) sent to the tissue via a silver mirror or through the FITC filter. Light intensity measured at the objective back aperture ranged from 1-25mW. Spatially-specific optogenetic experiments used a blue (473 nm) laser (Obis, Coherent) ranging from 0.6-2.7 mW measured at the back of the objective. Our preliminary uncaging experiments show the size of the laser spot to be <5 microns in diameter. However, in our optogenetic experiments the effect of the spot may be slightly larger (estimated at ~20-30 microns) due to the high light sensitivity of the CoChR rhodopsin. Optogenetic experiments were conducted in the presence of AP5 (50 μM) and either NBQX (5 μM) or CNQX (12.5 μM). Spatially-specific voltage clamp optogenetic experiments also included TTX (0.5 μM) and 4-AP (300 μM).

#### Two-photon calcium imaging

Calcium mas measured in SNc dopamine neuron dendrites and somas using the GCaMP6f mouse bred with the DAT-Cre mouse. All calcium imaging experiments were performed in the presence of AP5 (50 μM), NBQX (5 μM), and sulpiride (0.9 μM) to block NMDA-, AMPA- and dopamine D2-receptors. Two-photon calcium imaging was acquired on a custom microscope (Bruker). A Mai Tai Ti:sapphire laser (Spectra-Physics) was tuned to 980 nm. A 575 nm dichroic long-pass mirror was used to split the fluorescence signal through 607/45 nm and 525/70 nm filters (each notched at 470 nm) to above-stage and sub-stage multi-alkali photomultiplier tubes (Hamamatsu). Time-series spiral scans were acquired at 19-20 Hz. During acquisition, the blue optical stimulation laser was activated in 2 ms pulses at 19-20 Hz for 2 seconds. Optogenetic stimulation was synchronized with imaging frame rate to localize the light contamination from the laser to one area of the image. In a small fraction of cells, calcium signals were increased >110% of baseline during inhibition (17/463), presumably from light contamination or direct current activation from retrograde infection of ChR2. These cells were not included in analysis. Calcium signals were background-subtracted analyzed by manually-drawn regions of interest and processed with custom ImageJ macros and Igor (Wavemetrics) procedures.

#### Immunohistochemistry, clearing, confocal imaging, and neural reconstructions

After electrophysiology or imaging, slices were fixed overnight in 4% Paraformaldehyde (PFA) in phosphate buffer (PB, 0.1M, pH 7.6). Slices were subsequently stored in PB until immunostaining. CUBIC clearing was chosen because it does not quench fluorescence (Susaki et al., 2015). For the immunostaining/CUBIC clearing, all steps are performed at room temperature on shaker plate. Slices were placed in CUBIC reagent 1 for 1-2 days, washed in PB 3x 1 hour each, placed in blocking solution (0.5% fish gelatin (sigma) in PB) for 3 hours. Slices were directly placed in primary antibodies (sheep anti-TH and/or streptavidin Cy5 conjugate and/or rat anti-mCherry) at a concentration of 1:1000 in PB for 2-3 days. Slices were washed 3 times for 2 hours each and placed in secondary antibodies (Alexa 568 anti-sheep, or Alexa 488 anti-rat at 1:1000 in PB) for 2 days. After PB washed 3 times for 2 hours each, slices were placed in CUBIC reagent 2 overnight. Slices were mounted on slides in reagent 2 in frame-seal incubation chambers (Bio-Rad SLF0601) and coverslipped. Slices were imaged as tiled z-stacks on a Zeiss LSM 800 using Zen Blue software in the NINDS light imaging facility. Neural reconstructions were completed using these tiled z-stack images and were performed in Neurolucida (MBF bioscience). Additional neural reconstructions from two-photon images were conducted in NeuTube (Feng et al., 2015). Synaptophysin puncta density was determined by manually placed markers along each dendrite. Concentration of puncta with distance from the soma was determined by calculating the number of puncta in each Sholl ring (10 microns) and dividing it by the total dendritic length in that ring.

#### Drugs

Salts were purchased from Sigma. Alexa594 and Fluo5F (Life Technologies), 4-AP (Sigma, pH 7.34), TTX, gabazine, d-AP5, and NBQX (all purchased from Tocris) were prepared from aliquots stored in water. Sulpiride (Sigma) and CGP (Tocris) were dissolved in DMSO.

#### Computational model

All simulations were performed in Genesis simulation software (Bower and Beeman, 2007). A model of an SNc dopamine neuron was created as previously (Canavier et al., 2016; Tarfa et al., 2017). The morphology was based on a neural reconstruction and contained distinct SNr and SNc dendrites. This SNc neuron contains the following intrinsic channels: fast sodium (Tucker et al., 2012), leak sodium (Philippart and Khaliq, 2018), A-type potassium (Tarfa et al., 2017), Kv2 potassium (Khaliq and Bean, unpublished; Liu and Bean, 2014), Ih (Khaliq and Bean, unpublished, Migliore et al., 2008), and calcium channels L-type Cav1.3 and 1.2, N-type, R-type, and T-type Cav3.1 (Evans et al., 2013), and an SK channel (Evans et al., 2012; Hirschberg et al., 1998; Maylie et al., 2004). All channels are distributed at the soma and along all dendrites evenly, except Ih, which is located on dendrites >60 microns from the soma, CaT, which is located only on the dendrites, not the soma, and CaL1.3 which is located throughout the model neuron but at a higher concentration on the soma and proximal dendrites. The model was tuned to fire spontaneously at ~1-2 Hz. Inhibitory synaptic channels were modeled using the facsynchan object and included GABA-A (Galarreta and Hestrin, 1997) and GABA-B (Beckstead and Williams, 2007) receptor types. Depression and facilitation characteristics of the GABA-A receptor are based on our voltage-clamp recordings.

### Quantification and statistical analysis

Analysis was conducted in Igor (Wavemetrics). Mann-Whitney U tests were used to compare two samples. For multiple comparisons, Kruskal-Wallis tests determined significance of the dataset and post-hoc Mann-Whitney U tests determined significance between groups. Data in text is reported as Mean ± SEM and error bars on most graphs are ± SEM. Box plots show medians, 25 and 75% (boxes) confidence intervals, and 10 and 90% (whiskers) confidence intervals.

### Data and code availability

Computational model will be available on ModelDB and neural reconstructions will be made available on Neuromorpho.org upon publication of this manuscript.

### Key Resources Table

| REAGENT or RESOURCE | SOURCE | IDENTIFIER |
| --- | --- | --- |
| Antibodies |  |  |
| Sheep polyclonal anti tyrosine hydroxylase | Novus Biologicals | NB300-110 |
| Streptavidin cy5 conjugate | Invitrogen | SA1011 |
| Rat monoclonal anti mCherry (16D7) | Invitrogen | M11217<br>RRID:<br>AB_2536611 |
| Alexa 568 anti-sheep | Invitrogen | A-21099<br>RRID:<br>AB_2535753 |
| GFP anti-Rat | Invitrogen | A-21208<br>RRID:<br>AB_2535794 |
| **Bacterial and Virus Strains** |  |  |
| AAV1-hsyn-FLEX-CoChR-GFP | UNC vector core | Boyden, E. |
| AAV1-CAG-hChR2-mCherry.WPRE-SV40 | Addgene | 100054-AAV1 |
| AAV8.2-hEF1a-DIO-synaptophysin-mCherry | MIT Viral Gene Core | Neve, R.L. |
| **Chemicals, Peptides, and Recombinant Proteins** |  |  |
| SR95531 (Gabazine) | Tocris | Cat#1262 |
| CGP55845 | Tocris | Cat#1248 |
| D-AP5 | Tocris | Cat#0106 |
| CNQX | Tocris | Cat#1045 |
| NBQX | Tocris | Cat#1044 |
| Sulpiride | Sigma | Cat#8010 |
| Tetrodotoxin (TTX) | Tocris | Cat#1078 |
| 4-aminopyridine (4-AP) | Sigma | Cat#275875 |
| **Experimental Models: Organisms/Strains** |  |  |
| B6;129S-Pdyn(tm1.1(cre)/Mjkr)/LowIJ; (Pdyn-IRES-cre) | The Jackson Laboratory | Cat#027958 |
| B6;129S-Calb1(tm2.1(cre)/Hze)/J (Calb1-IRES2-cre-D) | The Jackson Laboratory | Cat#028532 |
| B6.129P2-Pvalb(tm1(cre)Abr)/J (PV-cre) | The Jackson Laboratory | Cat#017320 |
| B6;CBA-Tg(Lhx6-icre)1Kess/J (Lhx6-cre) | Nicoletta Kessaris |  |
| B6.SJL-Slc6a3(tm1.1(cre)Bkmn/J (DAT-IRES-cre) | The Jackson Laboratory | Cat#006660 |
| B6J.Cg-Gt(ROSA)26Sor(tm95.1(CAG-GCaMP6f)Hze)/MwarJ (Ai95(RCL-GCaMP6f)-D) | The Jackson Laboratory | Cat#028865 |

## Acknowledgements

Funding for this research was provided by an NINDS intramural research program grant NS003134 to Z.M.K. and NINDS/BRAIN Initiative K99NS112417 Career Development grant to R.C.E. We thank Dr. Carolyn Smith and the NINDS light imaging facility for assistance and use of the confocal microscopes.

