## Supplemental Figure S1 for "Functional dissection of basal ganglia inhibitory input onto SNc dopaminergic neurons"

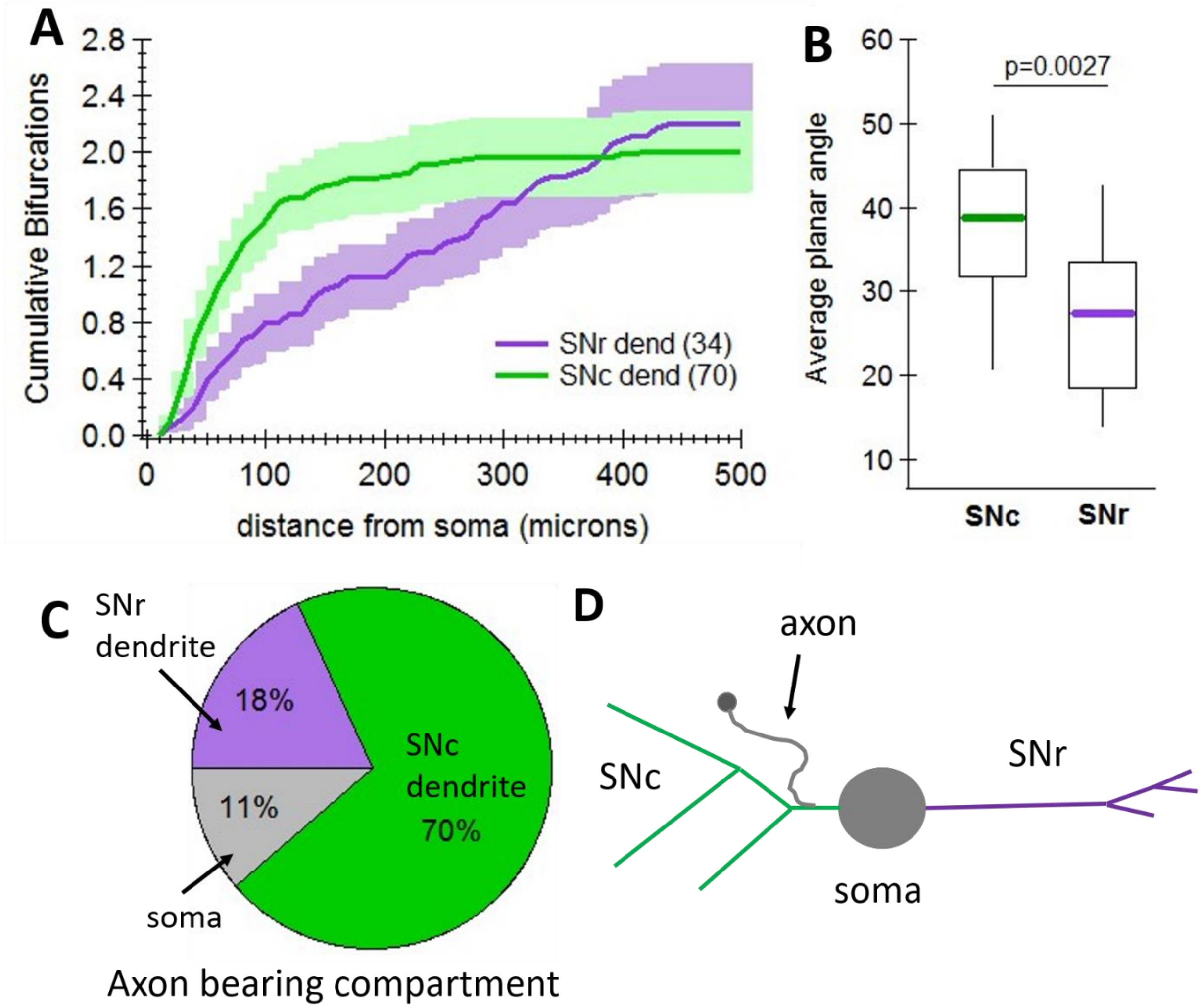


**Supplemental Figure S1. Distinct morphology of the SNr and SNc dendrites of SNc dopamine neurons. A.** SNc dendrites (n=70) branch more proximal to the soma than SNr dendrites (n=34). **B.** SNr dendrites branch at a narrower angle than SNc dendrites. **C.** The SNc dendrite is most often the axon bearing dendrite. **D.** Schematic of an SNc dopamine neuron showing the morphological characteristics for each dendrite type.
